# Meningeal-derived retinoic acid regulates neurogenesis via suppression of Notch and Sox2

**DOI:** 10.1101/2024.04.05.588348

**Authors:** Christina N. Como, Rebecca O’Rourke, Caitlin Winkler, Danae Mitchell, Luuli Tran, David Lorberbaum, Lori Sussel, Santos Franco, Julie Siegenthaler

**Author notes:** Corresponding Author <u>Contact Information:</u>, Julie Siegenthaler, PhD Associate Professor, University of Colorado, Anschutz Medical Campus Dept of Pediatrics, Section of Developmental Biology 12800 East 19th Ave, MS-8313, Aurora, CO 80045.

## Abstract

The meninges act as a regulator of brain development by secreting ligands that act on neural cells to regulate neurogenesis and neuronal migration. Meningeal-derived retinoic acid (RA) promotes neocortical neural progenitor cell cycle exit; however, the underlying molecular mechanism is unknown. Here, we used spatial transcriptomics and profiling of retinoic-acid receptor-α (RARα) DNA binding in *Foxc1*-mutant embryos that lack meninges-derived ligands to identify the neurogenic transcriptional mechanisms of RA signaling in telencephalic neural progenitors. We determined that meningeal-derived RA controls neurogenesis by suppressing progenitor self-renewal pathways Notch signaling and the transcription factor Sox2. We show that RARα binds in the *Sox2ot* promoter, a long non-coding RNA that regulates *Sox2* expression, and RA promotes *Sox2ot* expression in neocortical progenitors. Our findings elucidate a previously unknown mechanism of how meningeal RA coordinates neocortical development and insight into how defects in meningeal development may cause neurodevelopmental disorders.

## Introduction

Development of the central nervous system (CNS) requires precise intercellular crosstalk between neural and non-neural cells. One such example involves non-neural fibroblasts in the meninges. Meningeal fibroblasts surround the entire CNS during development and adulthood to regulate brain development and homeostasis^1,2^. The meninges are comprised of three layers (pia, arachnoid, dura) of fibroblasts and each layer can be identified by layer specific markers as early as embryonic day (E) 13 in mouse^3^. Meningeal fibroblasts help support brain development by secreting factors to regulate neural progenitor proliferation, neurogenesis, neuronal migration, neocortical neuron layering, and attachment of radial glial end feet^1,4–10^. Such factors include extracellular matrix proteins, Cxcl12, BMPs, Wnts and retinoic acid (RA)^5,7,10^. Despite identification of some meninges-derived factors, we have an incomplete understanding of the molecular pathways in neural cells that are downstream of meninges-produced signaling molecules.

Studies in mice and humans in which meningeal fibroblast secreted factors are reduced or absent have identified significant neurodevelopmental defects^7,11–17^, underscoring the importance of the meninges in brain development. *Foxc1* mutant mice are a well characterized model of aberrant meninges development. *Foxc1* mutant embryos have hypoplastic dorsal telencephalic meninges that lack layer-specific markers, including expression of meningeal factors like Cxcl12, BMPs, and RA that regulate neocortical development^7,16,18–20^. Meningeal defects in *Foxc1* mutant embryos cause lateral expansion of apical neural progenitors resulting in a strikingly lengthened neocortex and reduced numbers of neocortical neurons^7^. In humans, *FOXC1* point mutations cause cerebellar hypoplasia (Dandy-Walker malformation)^13,14^, hydrocephalus and white matter abnormalities.^21^ Furthermore, 6p25 deletion syndrome that includes a *FOXC1* gene deletion is characterized by neurodevelopmental delays and intellectual disability^13,22^, supporting a key role of *FOXC1* and the meninges in human brain development.

RA is a signaling molecule synthesized by meningeal fibroblasts during brain development and lacking in *Foxc1* mutant dorsal telencephalic meninges. Maternal supplementation of RA improves neurogenesis defects in *Foxc1* mutants, reducing neocortical length, increasing neural progenitor cell cycle exit and neuronal number^7^. This suggests meningeal-derived RA normally acts directly on neocortical neural progenitors to promote neurogenesis. RA functions by binding to DNA-bound retinoic acid receptors (RAR) to activate transcription of target genes^23,24^. RA-RARs are well known to promote neural progenitor differentiation by activating gene transcription^25–32^.

While these studies support that RA is an important factor regulating neurogenesis, the mechanism and transcriptional targets in neocortical neural progenitors are unknown. We utilized spatial transcriptomics of E14 *Foxc1* mutant mouse heads to study how absence of meningeal derived factors alters gene expression in telencephalic neural progenitors. This identified elevated Notch signaling components and Sox2 in *Foxc1* mutant telencephalic neural progenitors, pathways that repress neurogenesis^33,34^. Data from experiments using maternal RA supplementation, in utero electroporation and analysis of RARα binding in telencephalic neural progenitors support the model that RA, via binding to RARα, directly regulates expression of the long non-coding RNA *Sox2ot* to suppress Sox2 and downstream Notch signaling to promote neurogenesis. Collectively, this work provides previously unknown mechanistic insight into how meningeal-derived signals regulate neural progenitor self-renewal pathways.

## Results

### Spatial transcriptomics identifies pathways altered in neural progenitors of *Foxc1* mutants that fail to form normal forebrain meninges

To study how meningeal-derived factors like RA regulate neocortical development, we used mutant embryos that lack the transcription factor *Foxc1*, which is normally expressed by meningeal fibroblasts and blood vessels but not in neural cells (Supp. Fig. 1A). In E14 meninges above the neocortex, control meningeal fibroblasts expressed the fibroblast marker Collagen-1 (top) and the mature meningeal cellular markers CRABP2 (middle), and Raldh2 (bottom) that are important for RA signaling and synthesis (Supp. Fig. 1B, left). In the same region, *Foxc1^lacZ/lacZ^* (knockout or KO) meningeal fibroblasts expressed Collagen-1 but lacked expression of the RA synthesis proteins CRABP2 and Raldh2 (Supp Fig. 1B, right). In contrast to control meningeal fibroblasts, *Foxc1* mutant meningeal fibroblasts fail to differentiate and upregulate expression of RA synthesis enzymes *Aldh1a2* and *Rdh10* in the meninges between E12 and E14 (Supp. Fig. 1C, yellow and magenta arrows). This maturation failure in *Foxc1* mutants correlated with increased neocortical length at E13 and E14 (Supp. Fig. 1D, E). Elevated neocortical length results from increased neural progenitor self-renewal in *Foxc1* mutants^7,35,35^, supporting that defective meninges development underlies defects in anterior forebrain development.

**Figure 1.**
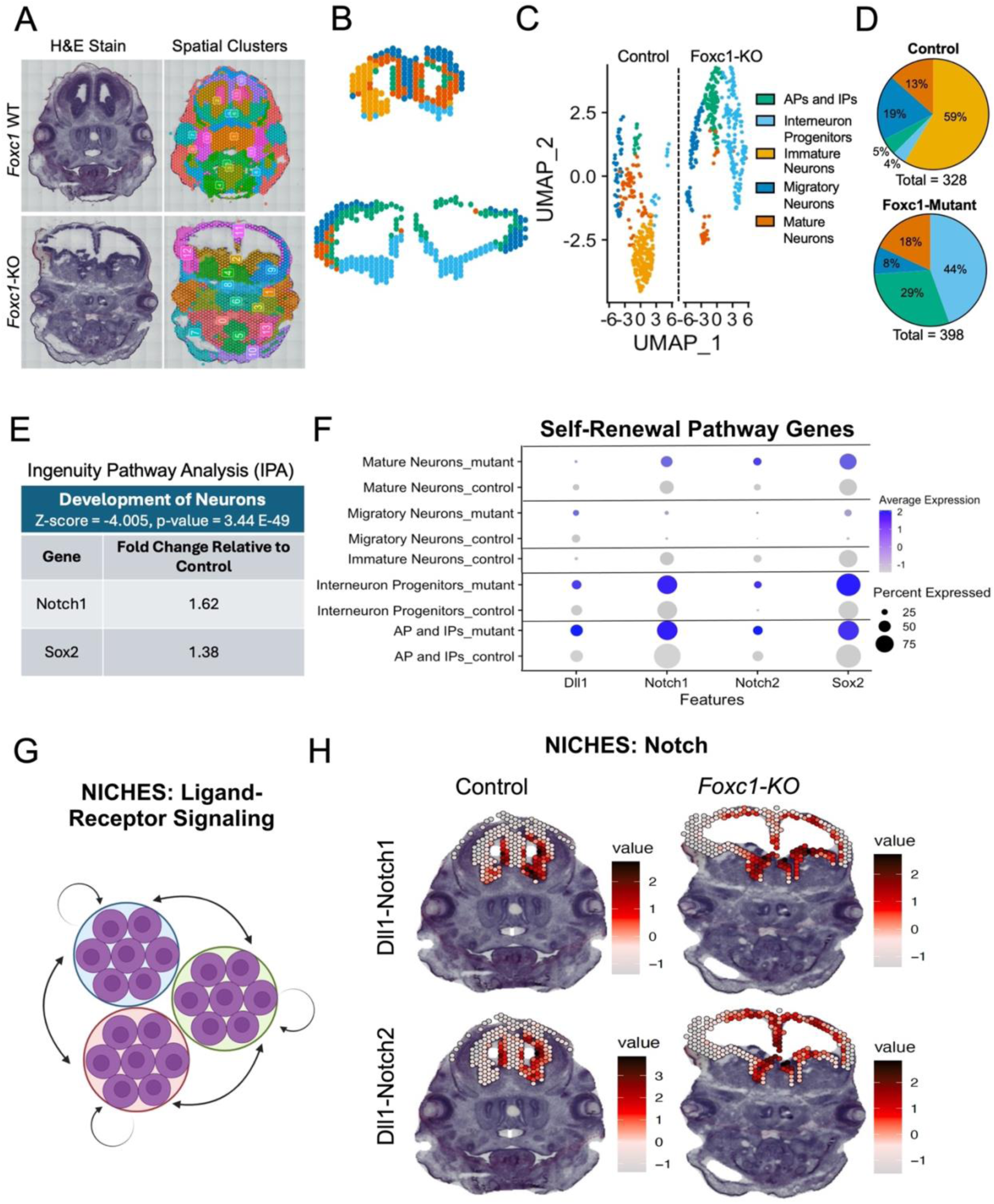
Spatial transcriptomic profiling of *Foxc1* mutants identifies increased self-renewal signaling pathway genes in neural progenitors. (A-C) Forebrain clusters from control and mutant tissue sections (A) were selected (B), and clusters were annotated and represented as a UMAP split by condition (Control or *Foxc1*-KO) (C). (D) The number of spots representing cellular populations were quantified for each condition. (E) Differential gene expression analysis between control and *Foxc1*-KOs using Ingenuity Pathway Analysis (IPA) showed the most significantly overlapped biological pathway “Development of Neurons” (z-score = −4.005, P-value = 3.44 E-49) and the predicted regulator *Notch1* and *Sox2*. (F) Dot plot depicting gene expression of *Dll1, Notch1, Notch2, and Sox2* in spatial clusters. (G-H) NICHES analysis was used to determine the differential ligand receptor signaling levels in spatial transcriptomic data (G) suggesting that Dll1-Notch1/2 signaling had stronger signaling interactions in *Foxc1*-KOs relative to controls in the cortical VZ and the VZ of the ganglionic eminences.

To identify pathways underlying meningeal control of anterior forebrain development, we applied 10x Visium Spatial Transcriptomics of whole heads to E14 control (Foxc1^+/+^, n=2) and *Foxc1*-KOs (n=2) (Fig. 1A). Spots corresponding to the forebrain were selected for both control and *Foxc1*-KOs and used for further analysis (Fig. 1B). Spatial transcriptomic cluster data was used to make a UMAP split by genotype and produced 5 different cellular clusters annotated as: Apical and Intermediate Progenitors (APs and IPs), Interneuron Progenitors, Immature Neurons, Migratory Neurons, and Mature Neurons (Fig. 1C). Cluster identities were annotated using markers reported in a E14 brain single cell atlas^36^ (AP and IPs: *Pax6*, *Eomes*; Interneuron Progenitors: *Dlx2*, *Sox2*; Immature Neurons: *Malat1*, *Lars2*; Migratory Neurons: *Cntn2*, *Neurod1*; Mature Neurons: *Tcf2*, *Neurod6*) (Supp. Fig. 2A, B). The number of spots representing cellular populations were graphed as a percentage of total spots per clusters for each genotype (Fig. 1D). Controls had a larger representation of neurons than progenitors (AP and IPs, 5%; Interneuron Progenitors, 4%; Immature Neurons, 59%, Migratory Neurons, 19%, Mature Neurons, 13%), whereas *Foxc1*-KOs had a larger representation of progenitors than neurons (AP and IPs, 29%; Interneuron Progenitors, 44%, Immature Neurons, 0%; Migratory Neurons, 8%; Mature Neurons, 18%). The decrease in neurons and increase in neural progenitors in *Foxc1*-KOs is consistent with prior analysis of *Foxc1* mutants showing increased progenitor self-renewal and fewer neocortical neurons^7^.

To identify potential pathways responsible for defective neurogenesis in *Foxc1* mutants, we performed differential gene expression analysis between control and *Foxc1*-KOs progenitor clusters using Ingenuity Pathway Analysis. This approach identified the most significantly overlapped biological pathway, the “Development of Neurons” (z-score = −4.005, P-value = 3.44 E-49), was significantly decreased, and two of the predicted regulators were *Notch1* and *Sox2* (increased 1.62-fold and 1.38-fold) (Fig. 1E). Corresponding downstream Notch targets were also found to be upregulated (*Hey1, Tp53, Hes1*, and *Hes5*). Consistent with this, *Foxc1*-KO samples had increased gene expression of Notch receptor *Dll1*, Notch ligands *Notch1*, *Notch2,* and *Sox2* across all spatial clusters (Fig. 1F). We used NICHES^37^ to model potential Notch ligand-receptor interactions within and between spatially adjacent spots (Fig. 1G) which suggested that Dll1-Notch1/2 had increased interactions in *Foxc1*-KOs relative to controls in the neocortical ventricular zone (VZ), which contains neural progenitors, and the VZ of the ganglionic eminences which contains interneuron progenitors (Fig. 1H). Collectively, this supports elevated expression of Notch pathway genes and *Sox2* in *Foxc1*-KO neural progenitor domains. The Notch pathway and Sox2 have documented functions in promoting neural progenitor self-renewal and regulating neurogenesis^33,34,38–41^, therefore this elevated expression likely underlies decreased neurogenesis in *Foxc1-KO* anterior forebrain regions.

### RA-RARα directly regulates expression of the long noncoding RNA Sox2ot, an inhibitor of Sox2

*Foxc1* mutants lack meningeal RA synthesis enzymes, have reduced neocortical RA levels and maternal RA diet supplementation improves defects in neocortical development in *Foxc1*-KO and *Foxc1* hypomorph (*hith*) mutants^7,42^. This implicates RA, through binding to RARs in neural progenitors, in regulation of self-renewal pathways Notch signaling and Sox2 to promote neurogenesis. To determine how RA might transcriptionally regulate expression of genes involved in neurogenesis identified in the spatial transcriptomic analysis, we first used a transcription factor DNA binding prediction algorithm (LASAGNA)^43^ to identify DNA binding sites for RA receptor RARα upstream of genes in the Notch pathway and *Sox2* (−1000 base pairs upstream to +200 base pairs downstream relative to transcription start sites). We opted to study RARα as it is the main receptor in neural progenitors^44,45^. We used *Rarb* as a positive control as it is known to be regulated by RA via direct binding of RARα^46–48^, this showed a high score for RARα binding, 94.08 (p-value = 2.5E-5, chromosomal location: chr14:16728769-16729905). Of the Notch pathway genes (*Notch1*, *Notch2, Hes1*, *Hes5, Hey1, Dll1*) and *Sox2* identified by spatial transcriptomics as upregulated in *Foxc1* mutant progenitors, only *Sox2* was predicted to have a RARα binding site with a similar prediction score to *Rarb* (score = 96.88, p value < 0, chromosomal location chr3:34649005-34650204). *Notch1, Notch2*, *Hes1, Hes5, and Dll1* had predicted binding sites, but a lower prediction score (Table 1). The chromosomal location of the predicted RARα binding site was upstream of *Sox2* but also within the long noncoding (lnc) RNA *Sox2ot* (Table 1). This was of interest as *Sox2ot* has previously been shown to inhibit *Sox2* transcription in neocortical progenitors^49,50^. We next analyzed published E14.5 cortical neural progenitor single nuclei ATAC-seq data^51^ to determine if predicted RARα binding sites are within regions of open chromatin, indicative of regions where proteins are bound to DNA and genes that are actively transcribed within neural progenitors. Predicted RARα binding sites were in ATAC peaks in *Sox2* and *Sox2ot*, notably near the transcriptional start sites (Fig. 2A, Table 1, yellow boxes), suggesting that these regulatory regions are potentially bound by RARα to regulate *Sox2* or *Sox2ot* transcription.

**Figure 2.**
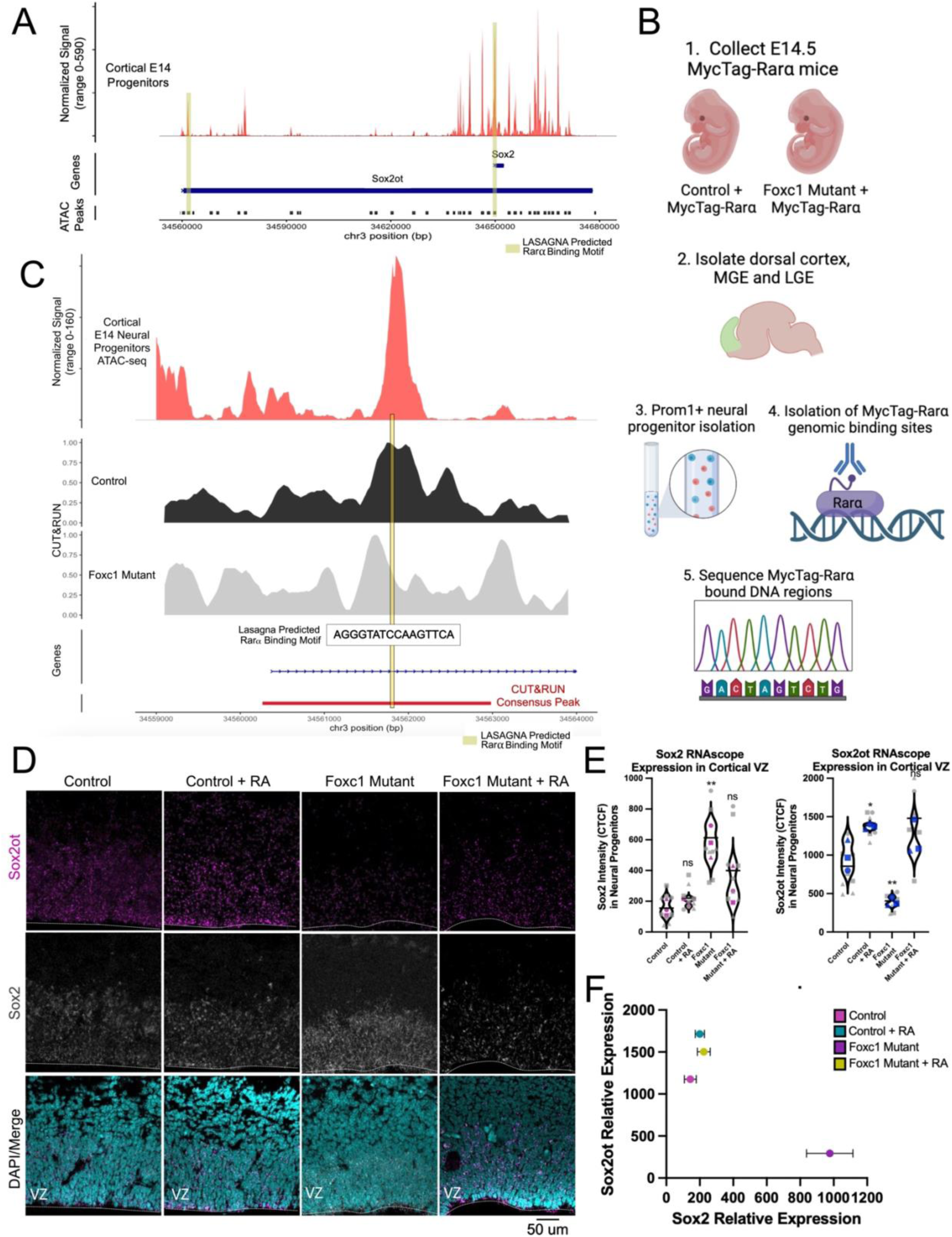
Retinoic acid receptor alpha (Rarα) regulates *Sox2* in neural progenitors by binding to *Sox2ot*. (A) E14 neocortical progenitor single nuclei ATAC-seq data depicting called peaks representative of open chromatin in the *Sox2ot*-*Sox2* gene locus. Predicted RARα binding motif location is shown in the promoter regions (yellow boxes). (B) Graphical depiction of neural progenitor cell isolation for Myc-RARα CUT&RUN to identify RARα-bound DNA in E14 neural progenitors. (C) Myc-RARα CUT&RUN and E14 neural progenitor snATAC-seq tracks depict location of consensus peak for RARα binding in control and *Foxc1* mutant, alignment with ATAC peak in the *Sox2ot* locus and site of RARα binding motif sequence. (D, E) Representative images (D) and CTCF quantification (E) of *Sox2* and *Sox2ot* expression in neocortical VZ. (F) Representation of inverse correlation between Sox2 and Sox2ot in controls and *Foxc1* mutants treated with or without RA. Means for each biological replicate (n = 3 embryos for each genotype-treatment) represented by colored shapes and corresponding technical replicates (n = 3 images per embryo) in grey circles, lines show mean. *, P < 0.05, **, P < 0.01. VZ = Ventricular Zone.

**Table 1.** RARα Predicted Binding Sites from Lasagna Motif Prediction.

| Gene | Score | p-value | Chromosome | CUT&RUN<br>Consensus Peak | Open Chromatin<br>Peak |
| --- | --- | --- | --- | --- | --- |
| *Rarb* | 94.08 | 2.5E-5 | chr14:16728769-16729905 | Yes | Yes |
| *Abtb2* | 88.06 | 5E-5 | chr2:103636080-103638179 | Yes | Yes |
| *Sox2* | 96.88 | 0 | chr3:34649005-34650204 | No | Yes |
| *Sox2ot* | 84.77 | 0.000125 | chr3:34555381-34561380 | Yes | Yes |
| *Notch1* | 10.39 | 0.00025 | chr2:26504822-26503623 | No | No |
| *Notch2* | 76.95 | 0.00055 | chr3:98012538-98013737 | Yes | Yes |
| *Hes1* | 74.23 | 0.00975 | chr16:30064357-30065556 | No | Yes |
| *Hes5* | 79.48 | 0.000275 | chr4:154959923-154961122 | No | Yes |
| *Dll1* | 8.49 | 0.0006 | chr17:15376823-15375624 | No | Yes |

To identify RARα DNA binding sites in telencephalic neural progenitors of control and *Foxc1* mutants, we crossed mice expressing an *Rara* allele in which the RARα protein is fused to a Myc sequence (MycTag-RARα) into our *Foxc1* mutant background (*Foxc1^hith/lacZ^*). We dissected the neocortices and ganglionic eminences from E14 *Foxc1* mutant and littermate controls expressing a single copy of the MycTag-RARα and dissociated tissue into single cell suspensions. We used magnetic bead separation to isolate Prom1+ neural progenitors and performed cleavage under targets and release using nuclease with (CUT&RUN) analysis with a Myc antibody to isolate and sequence RARα-bound DNA (Fig. 2B). Our analysis of MycTag-RARα bound sequences identified consensus peaks in known RARα target genes *Rarb* and *Atb2b2* that aligned with sites of open chromatin in these genes (Supp. Fig. 4A-B, Table 1). This supports the specificity of our approach to identify RARα-bound DNA. Analysis of MycTag-RARα bound sequences showed a called peak at the chromosomal region of *Sox2ot* in both control and *Foxc1* mutant samples (Fig. 2C, CUT&RUN Track). This aligned with the ATAC peak in the *Sox2ot* promoter region, containing the predicted RARα binding site (Fig. 2C, ATAC track). We did not identify consensus peaks in open chromatin regions near or within *Sox2* or Notch genes, except *Notch2* (Table 1). These data collectively support that RARα is bound to the *Sox2ot* locus in E14 neural progenitors.

We next tested if RA regulates *Sox2ot* and *Sox2* gene expression in neural progenitors using fluorescent in situ hybridization in sections from control and *Foxc1* mutants without treatment and following maternal RA supplementation, a treatment that improves neocortical lengthening in *Foxc1* mutants (Supp. Fig. 3). Compared to untreated controls, *Foxc1* mutants had significantly elevated *Sox2* in the neocortical VZ and significantly decreased expression of *Sox2ot* (Fig. 2D-E). RA treatment significantly increased VZ levels of *Sox2ot* in both control and *Foxc1* mutants as compared to the untreated conditions (Fig. 2D-E), restoring *Sox2* and *Sox2ot* expression in *Foxc1* mutants to control, untreated levels. This analysis revealed an inverse relationship of *Sox2ot* and *Sox2* expression in neural progenitors (Fig. 2F), supporting that RA, via RARα transcriptional activity, directly stimulates expression of *Sox2ot* to reduce expression of *Sox2*.

Spatial transcriptomics showed that *Sox2* expression and Notch ligands and receptors were all elevated in the interneuron progenitor cluster that represents the lateral ganglionic eminence (LGE) VZ. Further, *Foxc1* mutant LGE appeared qualitatively enlarged compared to controls and this appeared normalized with RA supplementation (Supp. Fig. 3C, yellow arrow). We observed similar changes in *Sox2* and *Sox2ot* expression in the LGE VZ (Supp. Fig. 4C-E). This suggests that LGE neural progenitors also rely on meningeal-derived RA to suppress Sox2.

### Maternal retinoic acid supplementation normalizes elevated Notch signaling and Sox2 in *Foxc1* mutant neural progenitors

Studies in mice in which Notch signaling is increased in neural progenitors identified similar phenotypes as *Foxc1* mutants, including lengthened neocortices, increased neural progenitor self-renewal, and decreased neurogenesis^52,53^. Sox2, a transcriptional regulator of Notch pathway genes^34,40^, is increased in *Foxc1* mutants and our data support that Sox2 is regulated by RA-RARα via *Sox2ot*. We therefore tested if RA supplementation normalizes expression of Notch pathway proteins and signaling in *Foxc1* mutants.

Cells in the neocortical VZ were analyzed for levels of Notch ligands and receptors via immunofluorescence followed by quantification using Calculated Total Cell Fluorescence (CTCF). Compared to untreated and RA-treated controls, *Foxc1* mutants had increased expression of Dll1 (yellow) and Notch1 (magenta) that were restored to control levels with RA treatment (Fig. 3A-C). To detect Notch signaling activity in neural progenitors, we combined immunofluorescence for Notch intracellular domain (NICD) (magenta) and Sox2 (yellow). Compared to untreated and RA treated controls, *Foxc1* mutants had increased levels of Sox2 and NICD and this was rescued with RA treatment (Figure 3D, E). Notch1, Dll1 and NICD were increased in the LGE of *Foxc1* mutants, and this was rescued with RA treatment (Supp. Fig. 4F-J). This supports that RA is upstream of Notch pathway proteins, potentially through direct regulation of *Sox2-Sox2ot*.

**Figure 3.**
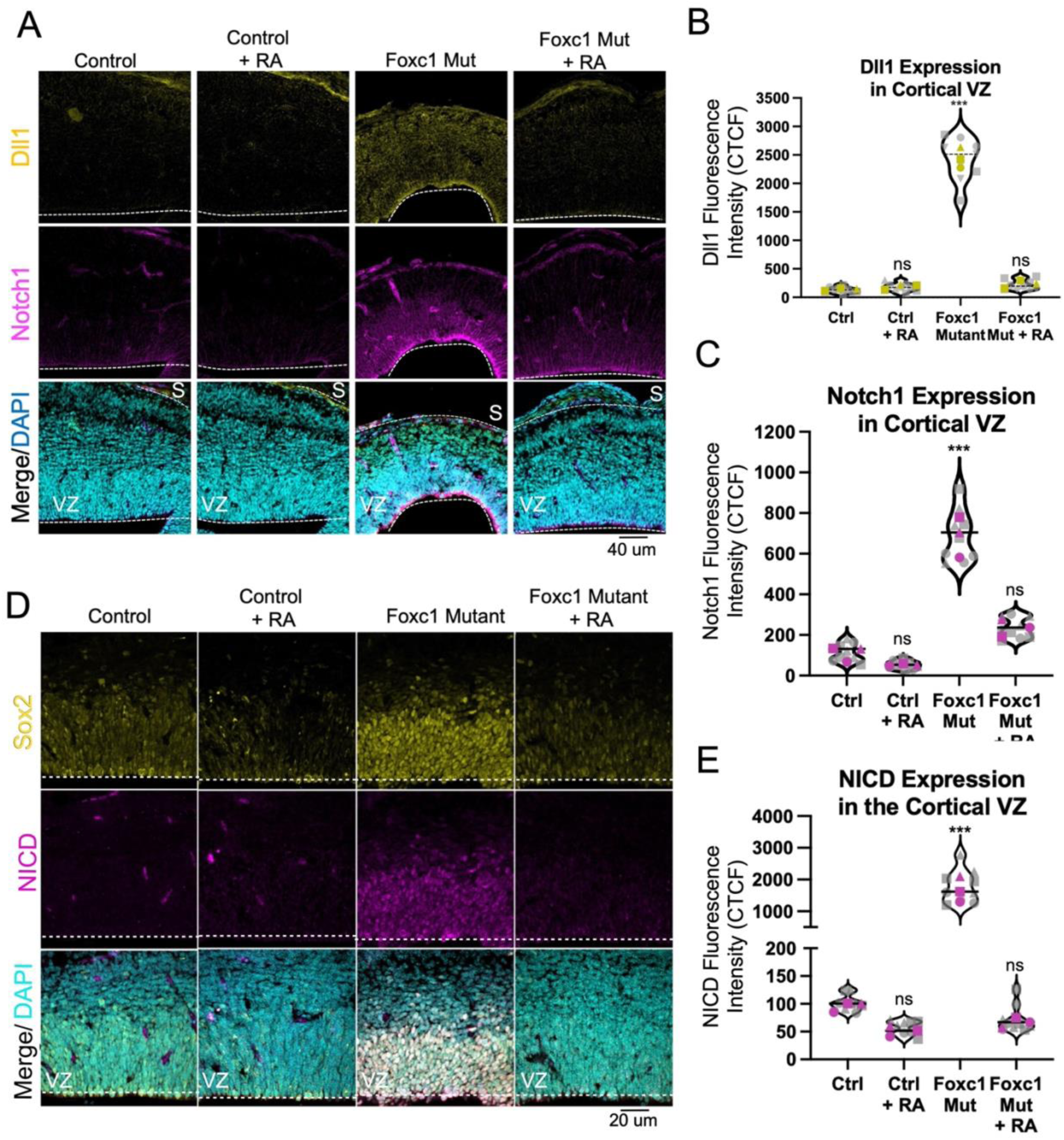
Elevated Notch signaling and Sox2 expression in Foxc1 mutants are rescued by RA treatment. (A-C) Representative images (A) and fluorescence intensity quantification (B, C) of Dll1 (yellow) and Notch1 (magenta) immunolabeling in control and *Foxc1* mutant neocortex exposed to control or RA supplemented diets. (D, E) Representative images (D) and fluorescence intensity quantification (E) of NICD (yellow) immunolabeling in control and *Foxc1* mutant neocortical VZ exposed to control or RA supplemented diets. Dotted lines outline the ventricular surface. For B, C, E, means for each biological replicate (n = 3 embryos for each genotype-treatment) represented by colored shapes and corresponding technical replicates (n = 3 images per embryo) in gray matching shapes, lines show mean. **, P < 0.01, ***, P < 0.001. VZ = Ventricular Zone, S = Skin.

We next analyzed how the appearance of meningeal defects, including absence of RA synthesis enzymes, correlated with neural progenitor expression of Sox2 protein and Notch signaling in *Foxc1* mutants. To do this we quantified Sox2 and NICD expression in the neocortex at timepoints that precede *Foxc1* mutant neocortical lengthening and defects in meningeal expression of RA synthesis enzymes (E12) and timepoints when these defects are apparent (E13, E14). Sox2 expression was similar in control and *Foxc1* mutant at E12 but significantly elevated at E13 and E14 (Supp. Fig. 1F). *Foxc1* mutant VZ expression of NICD at was not significantly different from control at E12 and E13 but significantly increased at E14 (Supp. Fig. 1G). Collectively, these data support a model in which meningeal-derived RA functions to regulate Sox2 expression and Notch signaling in VZ neural progenitors to promote neurogenesis in the neocortex.

### Inhibition of Notch signaling in neocortical progenitors increases neurogenesis in ***Foxc1* mutants.**

To test whether elevated Notch signaling in neocortical neural progenitors underlies increased self-renewal and decreased neurogenesis in *Foxc1* mutants, we used an in-utero electroporation (IUE) strategy to inhibit Notch signaling in neocortical progenitors in *Foxc1* mutant brains. We electroporated cells in the neocortical VZ with a GFP control plasmid or a plasmid containing dominant negative RBPJ (dnRBPJ) together with GFP (Fig. 4A, adapted from Tran et al., 2023^38^). RBPJ is the activating transcription factor that forms a transcriptional complex with NICD (elevated in *Foxc1* mutants) to positively regulate Notch target genes^38,54–56^. Control and *Foxc1* mutant (*Foxc1^hith/lacZ^*) embryos underwent IUE of a single hemisphere at E13.5 and were collected at E15.5. Control and *Foxc1* mutant heads at E15.5 showed intact brains and focal fluorescence representing the electroporation site (Fig. 4B, yellow arrow). Brains were sectioned on a vibratome and the part of the neocortex that contained GFP+ cells was quantified for expression of Hes1 as a readout of Notch signaling and knockdown via dnRBPJ electroporation (Fig. 4C). Relative to control brains electroporated with GFP, Hes1 signal was significantly increased in *Foxc1* mutant cells with control GFP electroporation but significantly decreased in control and *Foxc1* mutant cells electroporated with dnRBPJ plasmid electroporation (Fig. 4D). Consistent with our NICD analysis, this indicates that Notch signaling is increased in *Foxc1* mutants but can be reduced with expression of the dnRBPJ plasmid. The number of GFP+ cells were counted in the VZ (mostly Sox2+ progenitors), the intermediate zone (IZ), subventricular zone (SVZ) (mixture of progenitors and post-mitotic neurons) and cortical plate (CP, mostly post-mitotic neurons) and plotted to report the fraction of GFP+ cells in each region (Fig. 4E). Compared to control cells electroporated with GFP alone, the fraction of control cells electroporated with dnRBPJ in the VZ was significantly decreased, but the fraction in the CP was increased (Fig. 4F). This result supports inhibiting Notch signaling in control VZ cells at E13 promotes neurogenesis, consistent with a role of Notch signaling in progenitor cell maintenance and renewal^33,39,57^.

**Figure 4.**
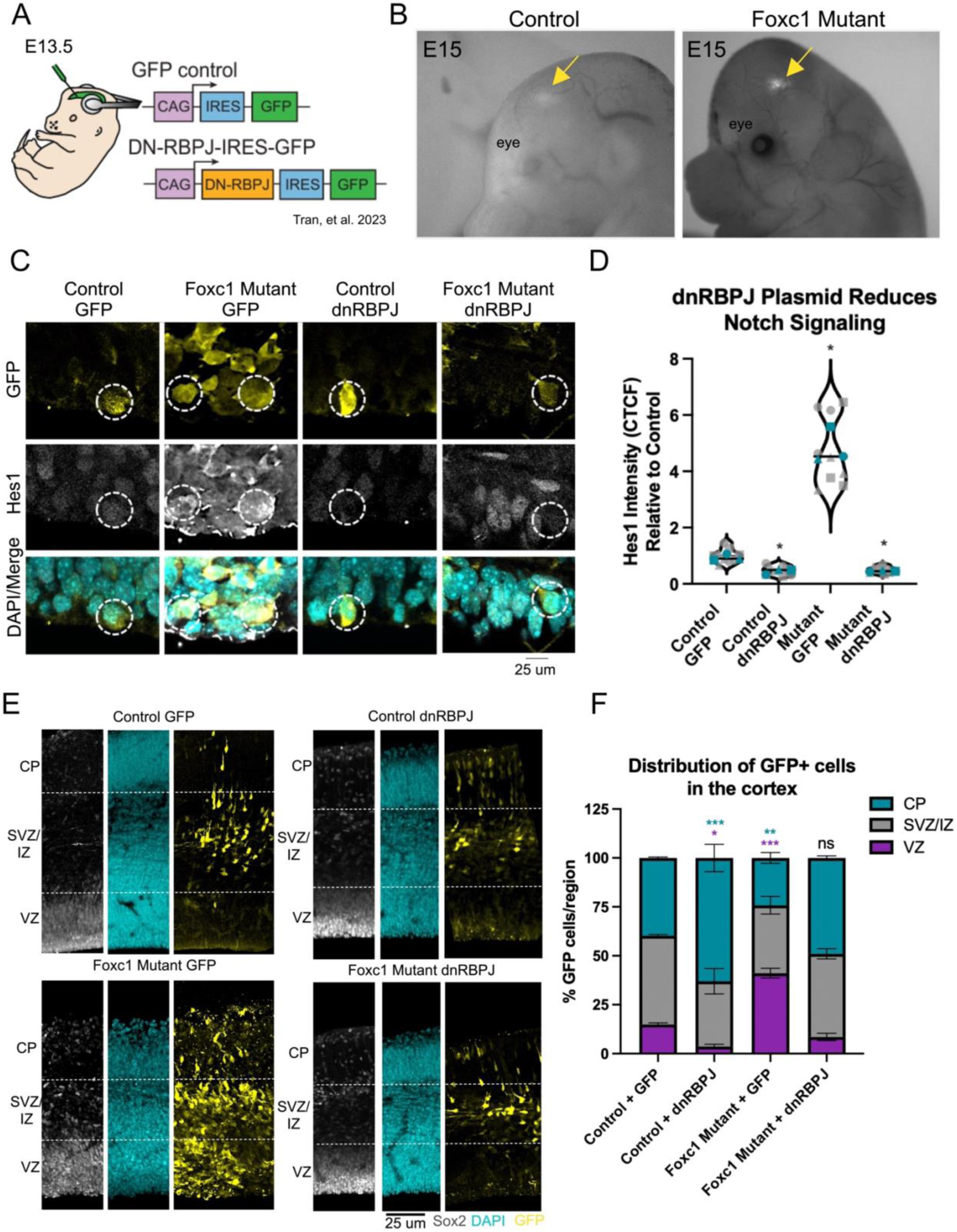
In utero electroporation of a Notch inhibitor rescues the *Foxc1* mutant neurogenesis phenotype. (A) Schematic of control GFP only plasmid and a dominant negative RBPJ (dnRBPJ) plasmid electroporated at E13.5 in control and *Foxc1* mutant ventricles and collected at E15.5. (B) Macroscopic images of whole embryos show fluorescence labeling (arrows) representing the IUE site in E15.5 control and *Foxc1* mutant neocortices. (C, D) Representative images and quantification of Hes1 expression intensity by CTCF in GFP+ cells in the VZ. Means for each biological replicate (n = 3 embryos for each genotype-treatment) represented by large colored shapes and corresponding technical replicates (n = 3 images per embryo) in small gray matching shapes, line and error bars show mean and SD. (E) Representative images of sections from control and *Foxc1* mutant neocortices electroporated with GFP plasmid only or dnRBPJ plasmid. DAPI and Sox2 co-labeling show CP and VZ, respectively and relative positioning of GFP+ cells in the different genotype-treatment conditions. (F) The number of GFP cells in the VZ, IZ/SVZ, CP were quantified and represented as a fraction of cells/region. *, P < 0.05, **, P < 0.01, ***, P < 0.001.

Compared to control cells electroporated with GFP, *Foxc1* mutant cells electroporated with GFP were significantly more often in the VZ and less frequently in the CP (Fig. 4F). This is consistent with increased neural progenitor cell self-renewal and reduced neuron production in *Foxc1* mutants^7^. In contrast, *Foxc1* mutant cells electroporated with dnRBPJ had a similar proportional distribution in the VZ and CP as control cells with GFP (Fig. 4F). There was no difference in the distribution of cells in the SVZ/IZ in all conditions. These data support that elevated Notch signaling in *Foxc1* mutant VZ cells underlies excessive progenitor self-renewal and decreased neurogenesis.

## DISCUSSION

Here, we provide evidence for a previously unknown mechanism of meningeal-derived RA in regulating neurogenesis. We show that during brain development, meningeal derived RA acts on neural progenitors in the neocortex and LGE to drive neurogenesis by suppressing self-renewal pathways Sox2 and Notch. Mechanistically, RA receptor RARα binds at the *Sox2ot-Sox2* locus in neural progenitors and RA binding drives increased *Sox2ot*, a lncRNA that suppresses *Sox2* in neural progenitors to promote neurogenesis^49,58^. Sox2 acts upstream of Notch to regulate the balance between neurogenesis and self-renewal^34,41,58,59^. These data support a model in which meningeal RA promotes neurogenesis in neural progenitors via RAR transcriptional regulation of *Sox2ot*, leading to decreased *Sox2* expression and downregulation of direct Sox2 targets, Notch ligand *Notch1* and receptor *Dll1* (**Graphical Abstract**).

The increase in neocortical progenitor self-renewal in *Foxc1* mutants is due to a lack of neural progenitor cell cycle exit, resulting in lateral expansion of the VZ and reduced neuron generation^7^. Our IUE experiment using dnRPBJ supports that this phenotype is due to increased Notch signaling, a potent stem cell renewal pathway. Our RA rescue experiments and the temporal correlation between meningeal defects in RA synthesis enzymes and neocortical lengthening collectively support that meningeal RA functions to reduce Notch signaling in both neocortical and LGE neural progenitors. This points to a previously unknown mechanism to how RA acts as a pro-neurogenic factor.

The increase in Notch signaling seen in *Foxc1* mutant neural progenitor cells was at the RNA and protein level, suggesting that RA plays a transcriptional role in regulating Notch signaling. However, we did not identify RARα binding sites near *Notch1* or *Dll1,* though we did at *Notch2*, indicating that RA-RAR regulation of Notch ligands and receptors is potentially indirect. *Notch1* is a direct transcriptional target of Sox2 in retinal neural progenitors^60^ and Sox2 is required for *Notch1* expression and progenitor maintenance during retinal development. Cell culture-based studies of Sox2 DNA binding in human neural progenitors (differentiated from embryonic stem cells lines) showed Sox2 binds to the *Dll1* promoter and regulates its expression^40^. This indicates excess Sox2 in *Foxc1* mutant neural progenitors stimulates elevated expression of Notch1, Dll1 and therefore Notch signaling. Alternatively, RA-RAR may normally inhibit the Notch pathway via non-genomic mechanisms. For example, we identified non-nuclear RARα inhibits Wnt-β-catenin signaling in brain endothelial cells via PKC-mediated degradation of β-catenin^61^.

We show RARα is bound upstream of *Sox2* within the promoter region of lncRNA *Sox2ot* and RA treatment stimulates expression of *Sox2ot,* supporting this lncRNA is a direct target of RA-RARα. RARs are typically constitutively bound to DNA and transcriptional activity is dependent on ligand binding. This constitutive binding may explain why RARα is bound at *Sox2ot* in *Foxc1* mutant neural progenitors. Lower *Sox2ot* expression in *Foxc1* mutants is likely due to lower RA levels in the *Foxc1* mutant neocortex^7^ that reduces RAR activity. LncRNAs such as *Sox2ot* can act as transcriptional regulators^62^. Two mechanisms of *Sox2ot* inhibition of *Sox2* transcription have been described. *Sox2* and *Sox2ot* expression is inversely correlated in mouse ESCs differentiated into NSCs; increased *Sox2ot* transcription reduces chromatin interaction between the *Sox2* promoter regions and *Sox2* super enhancer^50^. The inverse correlation in *Sox2-Sox2ot* mRNA mirrors our findings in neocortical and LGE neural progenitors. Further, *Sox2ot* in the neocortical neural progenitors negatively regulates Sox2 by directly interacting the transcriptional regulator YY1, which binds to the CpG island in the *Sox2* locus to suppress expression of *Sox2*^49^. Knockdown of *Sox2ot* in E13.5 neocortical progenitors via IUE increased Sox2 expression and cell proliferation as measured by BrdU incorporation^49^. This is like what we observe in *Foxc1* mutants that have significantly diminished *Sox2ot* in neural progenitors that is rescued with RA supplementation. This supports our model that meningeal-derived RA acts upstream to positively regulate *Sox2ot*, that in turn suppresses *Sox2* and downstream Notch signaling to promote neuron generation.

*Foxc1* mutant neocortical VZ phenotypes were also observed in the LGE VZ, including increased Notch signaling, Sox2 expression, and decreased *Sox2ot*, and expression was restored to control levels with RA supplementation. This suggests that meningeal-derived RA is also required for the progenitors in the VZ of the ganglionic eminences to correctly balance self-renewal and neuronal differentiation. We showed previously that ventral forebrain meninges, including expression of Raldh2, are much less affected in *Foxc1* mutants^7^. How might loss of dorsal forebrain meninges impact ventral forebrain neural progenitors? Cerebrospinal fluid (CSF) in the ventricles contain factors that act directly on VZ progenitors that line the ventricular surface to regulate proliferation and neurogenesis, examples include FGF2 and IGF. Studies in chick identified bioactive RA in CSF and provided evidence that the CSF is a route for RA to reach and act on the midbrain neuroepithelium^63,64^. More studies are needed to test the CSF as a route for meningeal-produced RA distribution and how this is impacted in *Foxc1* mutants with regional defects in meningeal development.

Although our study focuses on meningeal derived RA, other meningeal ligands could be involved, directly or indirectly, in modulation of Notch signaling or Sox2 expression. *Foxc1* mutants lack more than RA production from the dorsal forebrain meninges, other factors such as BMPs, Cxcl12, and Wnt ligands are also reduced. Meningeal Bmps^10^ have previously been implicated in neocortical development and meningeal Cxcl12^5^ is involved in multiple aspects of forebrain, midbrain and cerebellar development. Additionally, RA deficiency in *Foxc1* mutants reduces cerebral vasculature growth therefore disruption of neural progenitor-vessel association may decrease progenitor divisions^42,65^. Future work will identify the role of other meningeal-derived signals on regulation of self-renewal pathways like Notch and Sox2 and how these intersect with RA-RAR signaling.

## STAR METHODS

### Animals

Mice used for experiments were housed in specific pathogen-free facilities approved by AALAC and procedures were performed in accordance with animal protocols approved by the IACUC at The University of Colorado, Anschutz Medical Campus. The following mouse lines were used in this study: *Foxc1^LacZ/+^* mice and *Foxc1-hith* mutant alleles that have been described previously^11,66^, and a MycTag fused to RARα (description below). Timed matings between *Foxc1^LacZ/+^* mice were used to generate *Foxc1*-KO mice (*Foxc1^LacZ/LacZ^*) or a less severe mutant, *Foxc1*-mutants (*Foxc1^hith/LacZ^*). Mice were examined for a vaginal plug and the day the plug was found was E0.5. Littermate controls were used in all experiments, these were phenotypically normal genotypes *Foxc1^+/+^*, *Foxc1^LacZ/+^* _or_ *Foxc ^hith/+^*.

### Generation of MYC epitope tagged RARα mice

To knock-in a MYC epitope tag into the N terminus of RARα, we used homology directed repair (HDR) via CRISPR/Cas9 genomic engineering into B6N mice. Guide design was completed using CRISPOR and the Broad Institute sgRNA Design software, both of which have been refined to better identify off target events^67,68^. Results from each software were compared, and the guide which performed best using both algorithms was chosen. Guide activity was verified by incubating guide RNA and Cas9 protein with a PCR product containing the target sequence and comparing the ratio of cut to uncut PCR product.

Zygotes were cytoplasmically injected with guide RNA (TGGGCAGGAACTGCTATTGC TGG, Synthego at 5ng/ul), Cas9 (IDT, Alt-R® S.p. HiFi Cas9 Nuclease V3 [cat # 1081060] at 20ng/ul), and HDR DNA template (GCAaGAACTGCTgTTGCTtGCCAGATCCTCTTCTGAGATGAGTTTTTGTTCCATGCCAAGCAGCCACAGTCAGAAGAGCAGGCAAACAGTCTGGCAGGTAGTTGTGATGGCCT, from IDT dialyzed for final concentration of 25ng/ul) where appropriate. Injected zygotes were then transferred into pseudo pregnant recipients. 19 F0 pups were genotyped by PCR using primers outside the region to be modified followed by restriction enzyme digest. Of the pups born, 10 putative positive founders were identified. Several of these mice were bred to wildtype mice and F1 mice were sequenced using sanger sequencing to verify the precise mutation event. Intercrossing of sequence verified F1 mice within the same founder line resulted in homozygous knockin mice. These mice were healthy, bred normally, and had no observable deficits.

Genotyping protocol uses the 5’ primer (GTT GAC TAA CTT GGG ACT TGC) and 3’ primer (ACA ATG GTC ACC CTG GTC TAG) under standard thermocycling conditions with annealing temperature of 62C and extension time of 40sec for 36 cycles followed by digestion using Dde1 enzyme to cleave a unique site introduced during MYC epitope tag insertion. A wildtype allele yields a single 490bp product, while a mutant allele yields 193bp and 327bp products.

### Tissue processing, sectioning, immunostaining and in situ hybridization

Pregnant mice were anesthetized using isoflurane and embryos were collected at E12, E13, or E14 by removing the heads and placing them in 4% paraformaldehyde (PFA) overnight at 4 degrees C. Whole heads were placed into 20% sucrose overnight and embedded in OCT. Mouse heads were sectioned at 12 um increments using a cryostat and sections mounted on glass slides. Sections were blocked in 3% bovine serum albumin (BSA) + 0.05% Triton-X for 1 h at room temperature and incubated with primary antibodies at a dilution of 1:100. Following incubation with primary antibody(s), sections were incubated with appropriate Alexa Fluor-conjugated secondary antibodies and DAPI (Invitrogen, Carlsbad, CA, United States). In situ hybridization was completed using the ACD Bio Multiplex Fluorescent Detection assay kit (Neward, CA) and their probes against *Sox2* and *Sox2ot*.

### Spatial transcriptomics and analysis

To capture spatial transcriptomic data, we used the 10x Genomics Visium Platform following the manufacturer’s protocol for fresh frozen tissue from E14 control and *Foxc1*-KO whole mouse heads. Tissue was snap frozen in OCT using liquid nitrogen. Region matched cryosections of whole heads were collected via cryosection and directly mounted into the capture area (controls n=2, *Foxc1*-KO n=2). Hematoxylin stain and imaging, RNA capture, library prep and sequencing were performed by the Genomics and Microarray core at the University of Colorado, Anschutz Medical Campus. Samples were sequenced and data was packaged using the 10x Genomics SpaceRanger pipeline. Spatial data were integrated into the Seurat data pipeline for differential gene expression analysis. Differentially expressed genes were analyzed for pathway analysis using Ingenuity Pathway Analysis (IPA). Ligand-receptor analysis using NICHES^37^ was used to identify significant ligand-receptor signaling pathways between control and Foxc1-mutants. R Studio with version 2021.09.0, R version 4.3.1, and Seurat^69^ version 4.3.0.1 were used for analysis of Spatial Transcriptomic data.

### RA diet

Pregnant dams carrying control and *Foxc1^hith/Lacz^* embryos were fed an all-trans-retinoic acid (atRA) enriched diet (final concentration 0.35 mg/g food) consisted of atRA (Sigma-Aldrich) dissolved in corn oil and mixed with Bioserv Nutra-Gel Diet, Grain-Based Formula, Cherry Flavor. atRA diet was prepared fresh daily and provided *ad libitum* to pregnant wild-type females beginning in the afternoon of E10 through the day of collection on E14 as described previously^70^.

### In utero electroporation and injection

Pregnant dams carrying control and *Foxc1* mutant embryos were electroporated as described previously^71^. Survival surgeries were performed on timed pregnant mice (E13.5) to expose their uterine horns. Approximately 1 ul of endotoxin-free plasmid DNA was injected into an embryonic lateral ventricle. The concentrations of the injection solutions containing either DN-RBPJ-IRES-GFP or GFP control were as previously described^38^. Electroporation consisted of 5 pulses separated by 950 ms applied at 45 V. Embryos were put back into the abdominal cavity and were allowed to develop in utero and collected at E15.5. Embryonic brains were sectioned and processed as previously described^38^.

### CUT&RUN

To isolate neural progenitor cells, control and *Foxc1* mutant cortices and ganglionic eminences were dissected on ice, cut into fine pieces using forceps, and dissociated using a Papain kit (Worthington Biochemical; Freehold, New Jersey). Enrichment of apical progenitors was completed using the MACS cell isolation protocol magnetic Prom1 beads (Miltenyi, Bergisch Gladbach, Germany). To determine where RARɑ is genomically bound, we performed CUT&RUN analysis using the EpiCypher CUT&RUN assay (Durham, NC). Biological isotype antibody controls were used for all samples to determine background antibody binding. DNA library preparation and sequencing was performed by the Genomics and Microarray core at the University of Colorado, Anschutz Medical Campus. Data was aligned using the nf-core CUT&RUN analysis pipeline^72^ with the following parameters: Spike-in normalization, SEACR as the peak caller, ‘norm’ normalization, relaxed threshold, and consensus peak requirement of 3 replicates. Plots were produced using Signac^73^.

### snATAC seq analysis

Publicly available snATACseq data from E14.5 cortical neural progenitors was downloaded from GEO (GSM5091383), converted to fastq files, and generated an h5 matrix file and fragments file with cellranger_atac count, summits were visualized using Integrated Genome Viewer and objects were generate as a Seurat object using Signac^73^.

### Microscopy and image analysis

Confocal images were obtained using a Zeiss 900 Laser Scanning Microscope with associated Zeiss Zen Blue software. Images were processed in FIJI Image J and Graphic Software. For quantification of protein and in situ fluorescence intensity, calculated total cell fluorescence (CTCF) was used for individual cells within the VZ for Dll1, Notch1, NICD, Sox2, Hes1, and *Sox2ot*. The number of GFP+ cells within the VZ and CP post electroporation was determined by counting individual nuclei within the defined regions as determined by DAPI and Sox2.

### Statistical Analysis

Statistical analyses were performed using GraphPad Prism, where two-way ANOVAs and multiple comparisons were performed in instances with more than two groups.

## Data Availability

Upload to GEO pending.

## Acknowledgments

The authors thank the leadership and staff of the Genomics and Microarray Core at the CU Anschutz Medical Campus for expert processing of samples and sequencing services and the Regional Mouse Genetics Core Facility at National Jewish for generation of the MycTag-RARα mouse line. The authors would like to thank the members of the Siegenthaler Lab and members of CC’s thesis committee (Dr. Emily Bates, Dr. Kristin Artinger, Dr. Joe Brzezinski, and Dr. Mark Dell’Acqua) for fruitful discussions and feedback that helped shape this project. This work was supported by funding from the NIH, specifically NINDS (F31NS130941 to CC, R01 NS098273 to JAS, R01 NS124166 to SJF, F31 NS129289 to LNT) and from NIDDK R01 DK118155 (LS), R00 DK128537 and P30 DK020572 (DL). Pilot Award for the spatial transcriptomics was provided from the RNA Biosciences Initiative at CU Anschutz. All schematics in the figures were made with BioRender.

## KEY RESOURCES TABLE

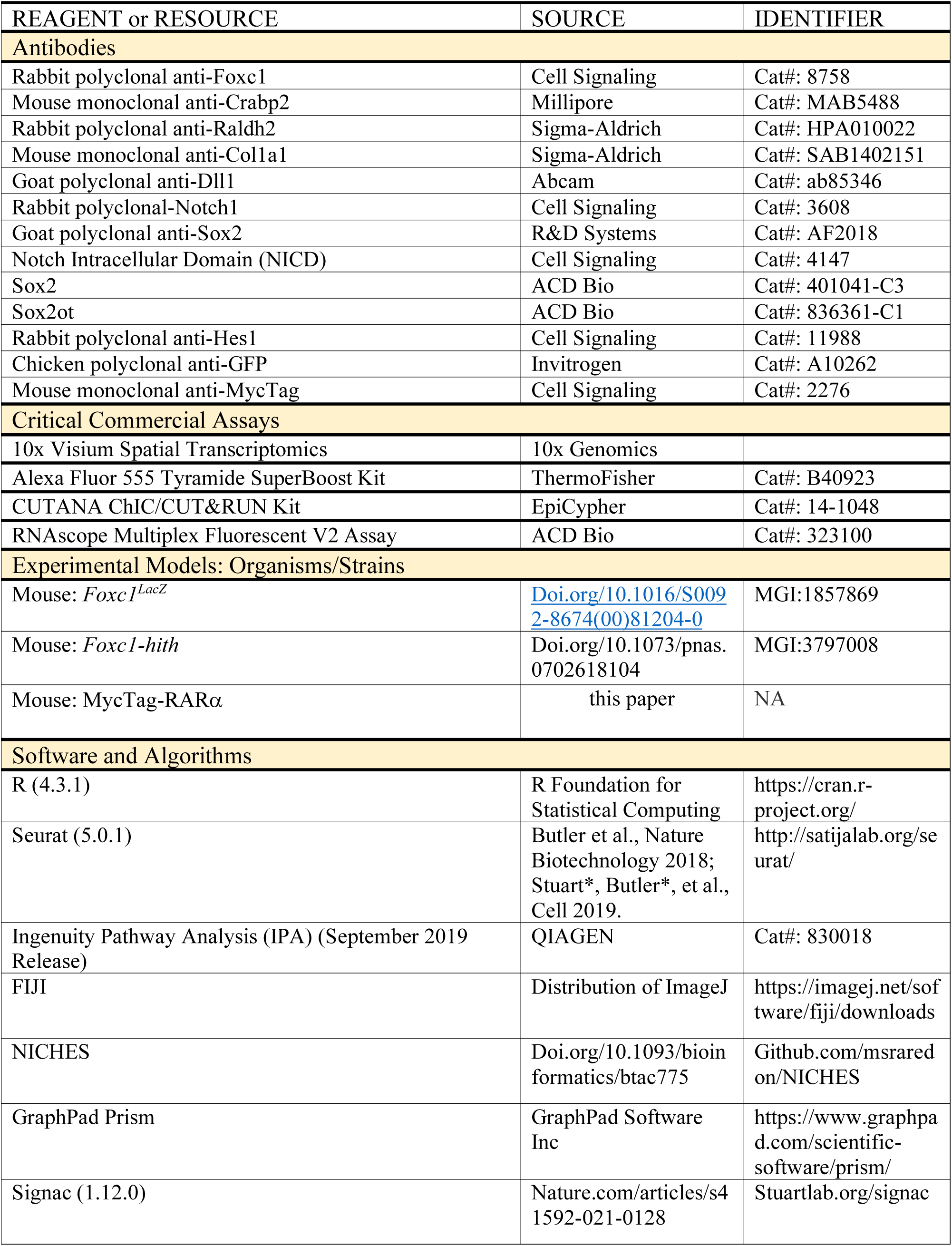
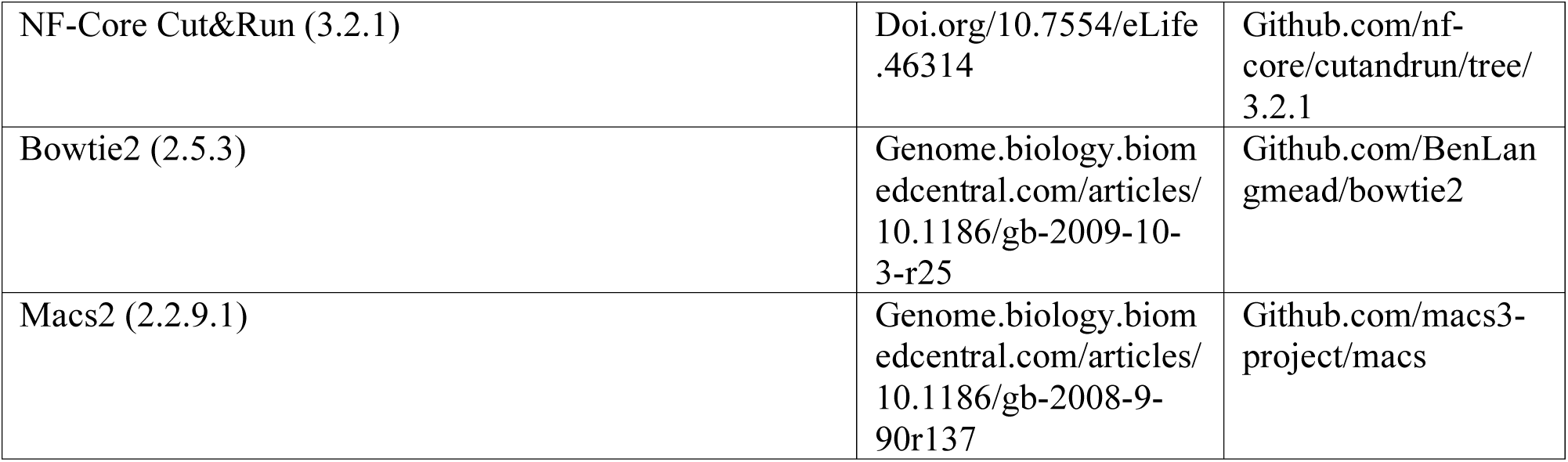

| REAGENT or RESOURCE | SOURCE | IDENTIFIER |
| --- | --- | --- |
| **Antibodies** |  |  |
| Rabbit polyclonal anti-Foxc1 | Cell Signaling | Cat#: 8758 |
| Mouse monoclonal anti-Crabp2 | Millipore | Cat#: MAB5488 |
| Rabbit polyclonal anti-Raldh2 | Sigma-Aldrich | Cat#: HPA010022 |
| Mouse monoclonal anti-Col1a1 | Sigma-Aldrich | Cat#: SAB1402151 |
| Goat polyclonal anti-Dll1 | Abcam | Cat#: ab85346 |
| Rabbit polyclonal-Notch1 | Cell Signaling | Cat#: 3608 |
| Goat polyclonal anti-Sox2 | R&D Systems | Cat#: AF2018 |
| Notch Intracellular Domain (NICD) | Cell Signaling | Cat#: 4147 |
| Sox2 | ACD Bio | Cat#: 401041-C3 |
| Sox2ot | ACD Bio | Cat#: 836361-C1 |
| Rabbit polyclonal anti-Hes1 | Cell Signaling | Cat#: 11988 |
| Chicken polyclonal anti-GFP | Invitrogen | Cat#: A10262 |
| Mouse monoclonal anti-MycTag | Cell Signaling | Cat#: 2276 |
| **Critical Commercial Assays** |  |  |
| 10x Visium Spatial Transcriptomics | 10x Genomics |  |
| Alexa Fluor 555 Tyramide SuperBoost Kit | ThermoFisher | Cat#: B40923 |
| CUTANA ChIC/CUT&RUN Kit | EpiCypher | Cat#: 14-1048 |
| RNAscope Multiplex Fluorescent V2 Assay | ACD Bio | Cat#: 323100 |
| **Experimental Models: Organisms/Strains** |  |  |
| Mouse: *Foxc1*<sup>LacZ</sup> | [Doi.org/10.1016/S0092-8674(00)81204-0](https://doi.org/10.1016/S0092-8674%2800%2981204-0) | MGI:1857869 |
| Mouse: *Foxc1-hith* | Doi.org/10.1073/pnas.0702618104 | MGI:3797008 |
| Mouse: MycTag-RARα | this paper | NA |
| **Software and Algorithms** |  |  |
| R (4.3.1) | R Foundation for Statistical Computing | https://cran.r-project.org/ |
| Seurat (5.0.1) | Butler et al., Nature Biotechnology 2018; Stuart*, Butler*, et al., Cell 2019. | http://satijalab.org/seurat/ |
| Ingenuity Pathway Analysis (IPA) (September 2019 Release) | QIAGEN | Cat#: 830018 |
| FIJI | Distribution of ImageJ | https://imagej.net/software/fiji/downloads |
| NICHES | Doi.org/10.1093/bioinformatics/btac775 | Github.com/msraredon/NICHES |
| GraphPad Prism | GraphPad Software Inc | https://www.graphpad.com/scientific-software/prism/ |
| Signac (1.12.0) | Nature.com/articles/s41592-021-0128 | Stuartlab.org/signac |
| NF-Core Cut&Run (3.2.1) | [Doi.org/10.7554/eLife.46314](https://doi.org/10.7554/eLife.46314) | [Github.com/nf-core/cutandrun/tree/3.2.1](https://github.com/nf-core/cutandrun/tree/3.2.1) |
| Bowtie2 (2.5.3) | [Genome.biology.biomedcentral.com/articles/10.1186/gb-2009-10-3-r25](https://genome.biology.biomedcentral.com/articles/10.1186/gb-2009-10-3-r25) | [Github.com/BenLangmead/bowtie2](https://github.com/BenLangmead/bowtie2) |
| Macs2 (2.2.9.1) | [Genome.biology.biomedcentral.com/articles/10.1186/gb-2008-9-90r137](https://genome.biology.biomedcentral.com/articles/10.1186/gb-2008-9-90r137) | [Github.com/macs3-project/macs](https://github.com/macs3-project/macs) |

