## Supplemental Data for "Meningeal-derived retinoic acid regulates neurogenesis via suppression of Notch and Sox2"

### Graphical Abstract

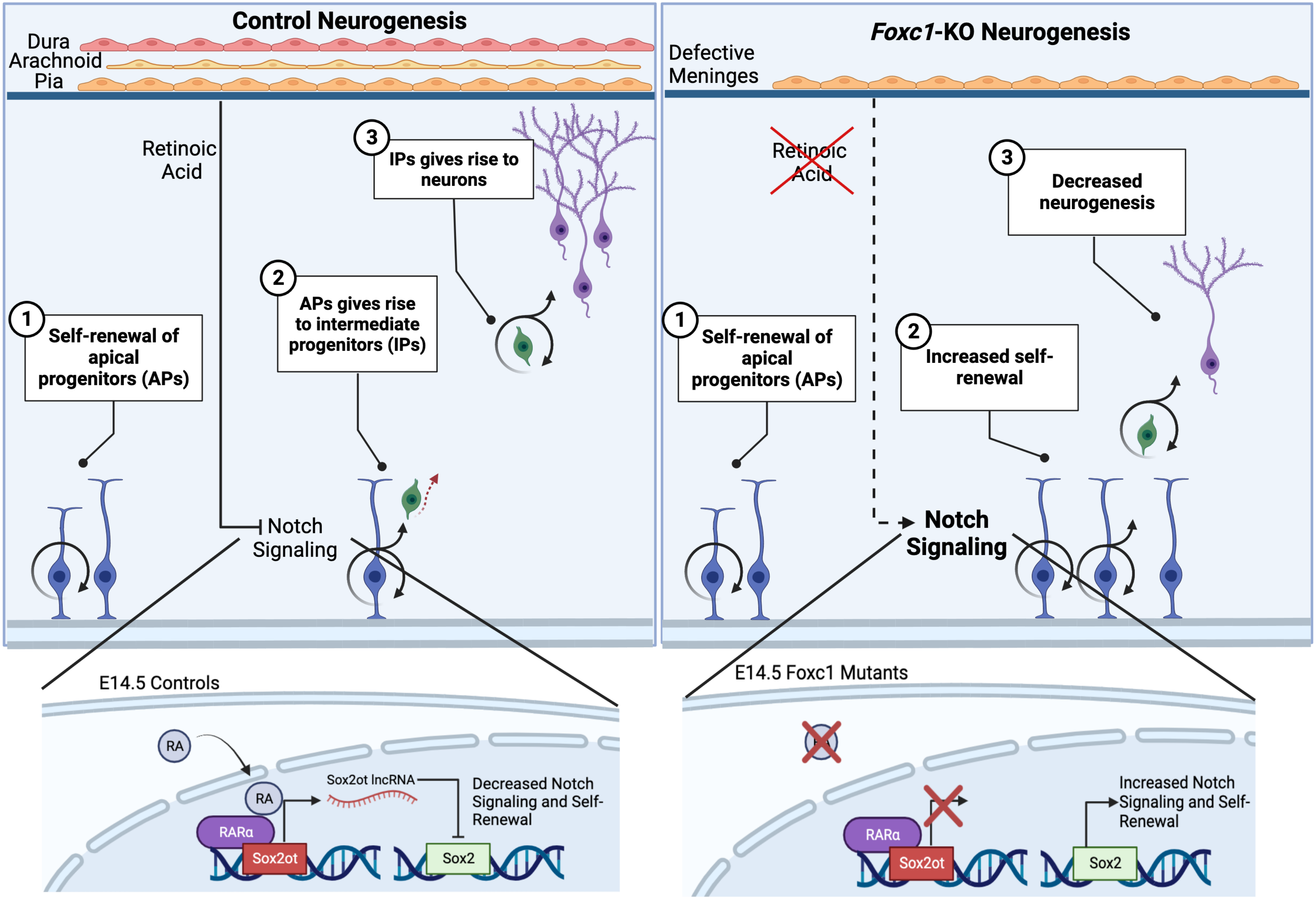

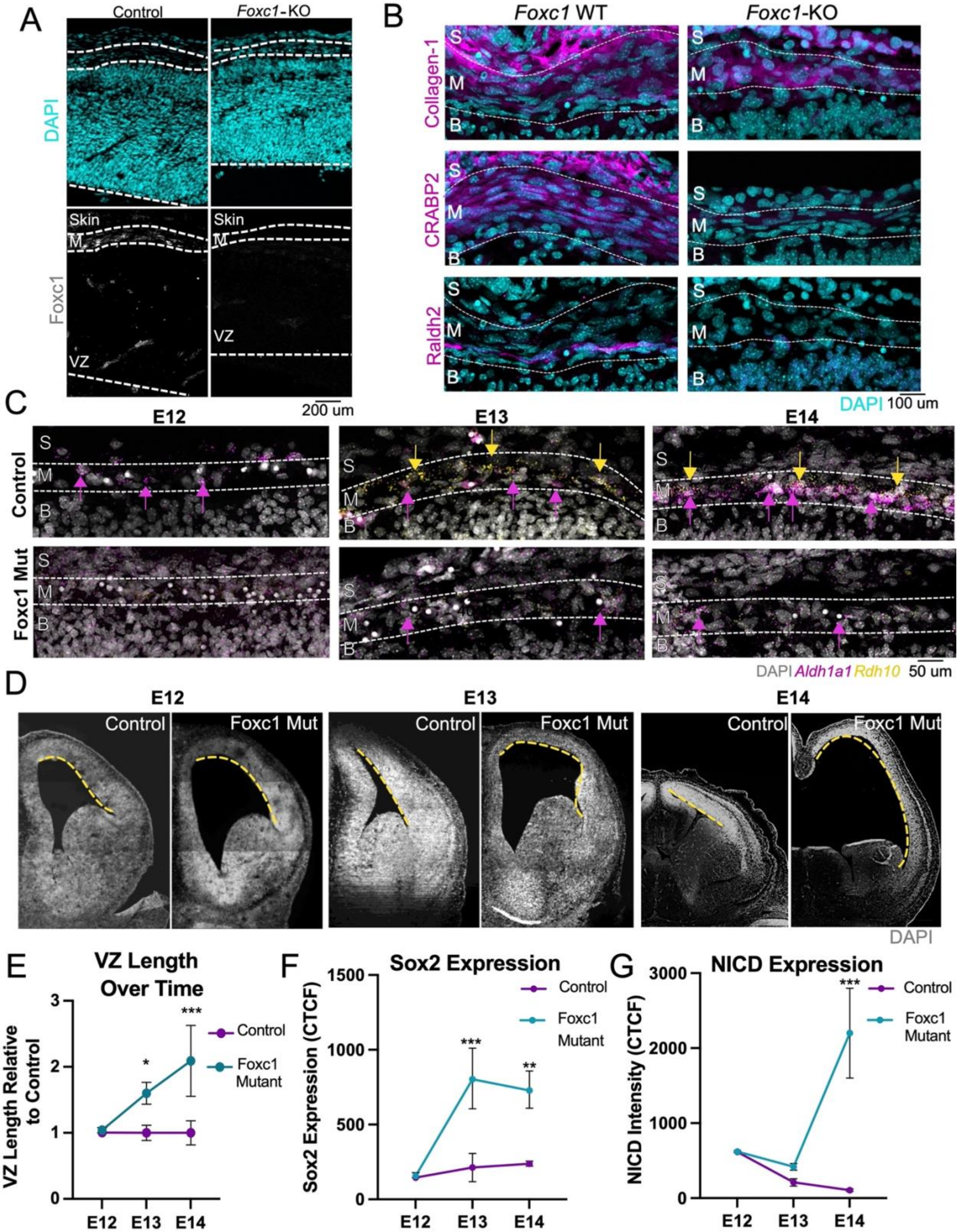
A
Control
Foxc1-KO
DAPI
Skin
M
Foxc1
VZ
200 um
B
Foxc1 WT
Foxc1-KO
Collagen-1
CRABP2
Raldh2
S
M
B
DAPI 100 um
C
E12
E13
E14
Control
Foxc1 Mut
DAPI Aldh1a1 Rdh10 50 um
D
Control
Foxc1 Mut
DAPI
E
VZ Length Over Time
VZ Length Relative to Control
Control
Foxc1 Mutant
F
Sox2 Expression
Sox2 Expression (CTCF)
G
NICD Expression
NICD Intensity (CTCF)

**Supplementary Figure 1. Appearance of defective neocortical development in *Foxc1* mutants correlates with timing of defects in meninges development.** Foxc1 is expressed in the meninges and not in neocortex, besides blood vessels, in control (A, left; grey), and *Foxc1*-KOs lack Foxc1 protein expression (A, right). Immunofluorescence shows control meningeal fibroblasts express Collagen-1 (top), CRABP2 (middle), and Raldh2 (bottom). *Foxc1*-KO meningeal fibroblasts express Colllagen-1 but not the mature cellular markers CRABP2 or Raldh2 (B). RNAscope for RA synthesizing enzymes *Aldh1a2* (encodes Raldh2) (magenta) and *Rdh10* (yellow) at E12, E13, and E14 show low expression at E12, and an increase at E13 to E14 in controls, with little to no signal in *Foxc1* mutants across timepoints (C). (D, E) Qualitative images and quantification of the ventricular zone length (VZ) in control and *Foxc1* mutants at E12, E13, and E14. (F, G) CTCF analysis of NICD and Sox2 protein expression in control and *Foxc1* mutants at E12, E13, and E14. S – Skin; M – Meninges; B – Brain. *, $P < 0.05$, **, $P < 0.01$, ***, $P < 0.001$.

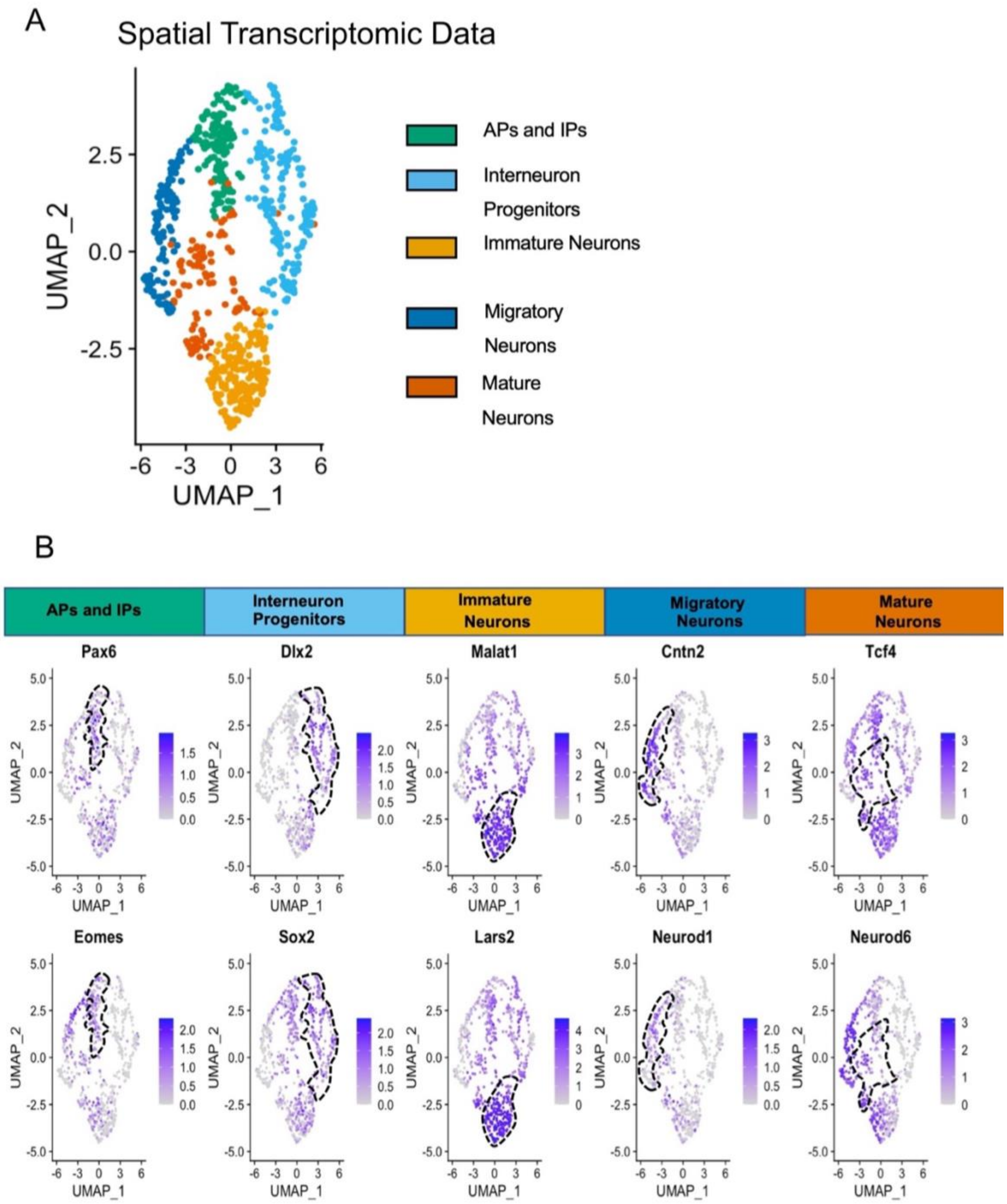


**Supplementary Figure 2. Spatial transcriptomics cluster annotation.** (A, B) Five spatial clusters were annotated based on differential gene expression: AP and IPs (*Pax6, Eomes*), Interneuron Progenitors (*Dlx2, Sox2*), Immature Neurons (*Malat1, Lars2*), Migratory Neurons (*Cntn2, Neurod1*), and Mature Neurons (*Tcf4, Neurod6*).

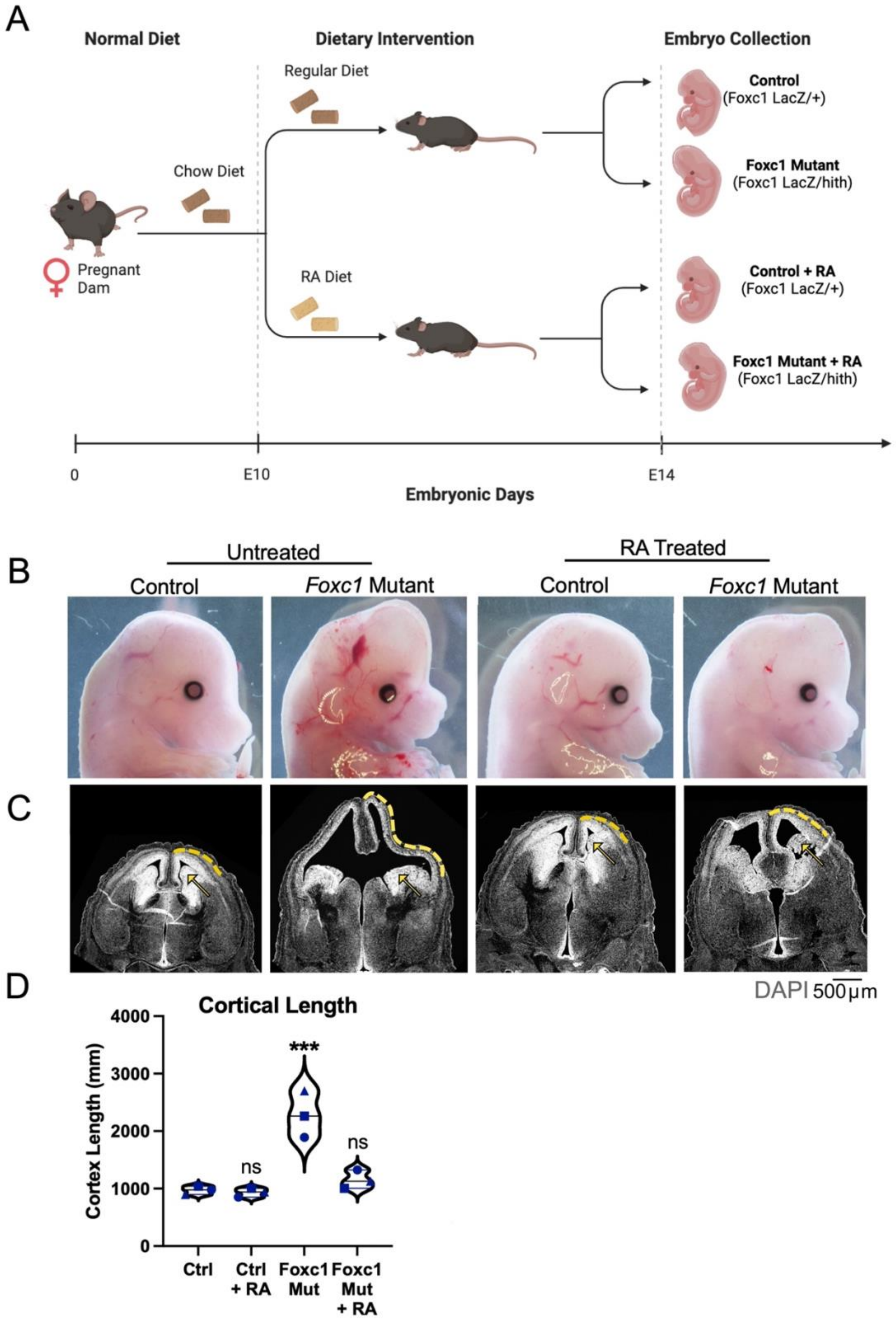


**Supplementary Figure 3. Maternal RA supplementation rescues the increased cortical length in E14 Foxc1 mutants.** (A) Graphical depiction of RA feeding paradigm where pregnant dams at E10 were either fed a control or RA supplemented diet and embryos were collected at E14. (B, C) Macroscopic images of whole E14 embryos (B) and microscopic images of whole head sections (C) of control and *Foxc1* mutants exposed to control of RA diet show improvement in cerebral hemisphere size and neocortical lengthening in *Foxc1* mutant with RA diet. (D) Quantification of the neocortical length (yellow dotted line along cortex, C). ***, P < 0.001.

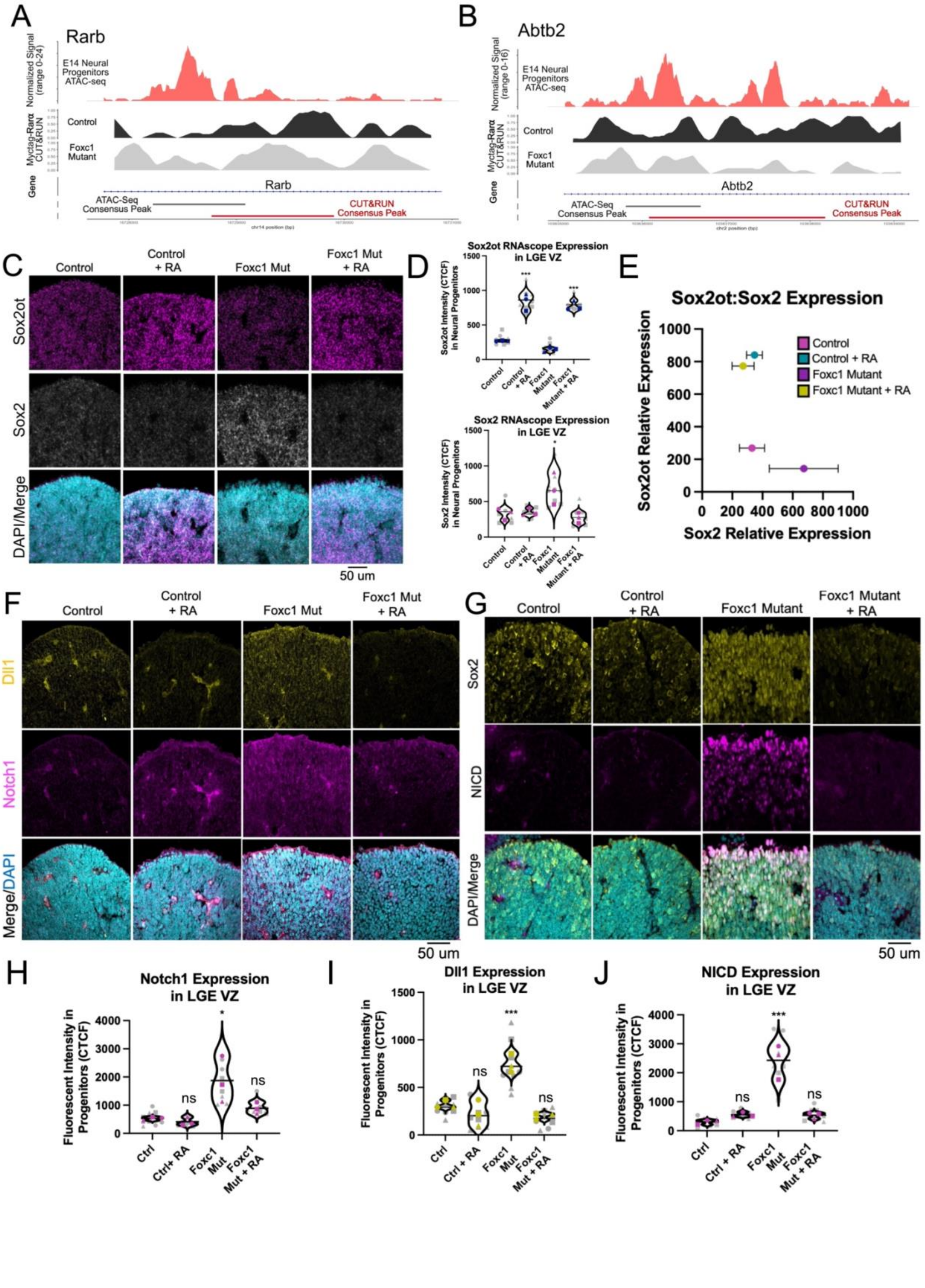
A
Rarb
Normalized Signal (range 0-24)
E14 Neural Progenitors ATAC-seq
Myctag-Rarα CUT&RUN
Control
Foxc1 Mutant
Gene
Rarb
ATAC-Seq Consensus Peak
CUT&RUN Consensus Peak
B
Abtb2
Normalized Signal (range 0-16)
E14 Neural Progenitors ATAC-seq
Myctag-Rarα CUT&RUN
Control
Foxc1 Mutant
Gene
Abtb2
ATAC-Seq Consensus Peak
CUT&RUN Consensus Peak
C
Control
Control + RA
Foxc1 Mut
Foxc1 Mut + RA
Sox2ot
Sox2
DAPI/Merge
50 um
D
Sox2ot RNAscope Expression in LGE VZ
Sox2ot Intensity (CTCF) in Neural Progenitors
Control
Control + RA
Foxc1 Mutant
Foxc1 Mutant + RA
Sox2 RNAscope Expression in LGE VZ
Sox2 Intensity (CTCF) in Neural Progenitors
E
Sox2ot:Sox2 Expression
Control
Control + RA
Foxc1 Mutant
Foxc1 Mutant + RA
Sox2ot Relative Expression
Sox2 Relative Expression
F
Control
Control + RA
Foxc1 Mut
Foxc1 Mut + RA
Dll1
Notch1
Merge/DAPI
50 um
G
Control
Control + RA
Foxc1 Mutant
Foxc1 Mutant + RA
Sox2
NICD
DAPI/Merge
50 um
H
Notch1 Expression in LGE VZ
Fluorescent Intensity in Progenitors (CTCF)
Ctrl
Ctrl+ RA
Foxc1 Mut
Foxc1 Mut + RA
I
Dll1 Expression in LGE VZ
Fluorescent Intensity in Progenitors (CTCF)
Ctrl
Ctrl + RA
Foxc1 Mut
Foxc1 Mut + RA
J
NICD Expression in LGE VZ
Fluorescent Intensity in Progenitors (CTCF)
Ctrl
Ctrl + RA
Foxc1 Mut
Foxc1 Mut + RA

861

**Supplementary Figure 4. Myc-RAR$\alpha$ consensus peaks in known RAR transcriptional targets and *Foxc1* mutants have increased Notch signaling in the LGE VZ.** (A, B) Myc-RAR$\alpha$ CUT&RUN consensus peaks were identified in control and *Foxc1* mutant with corresponding open chromatin ATAC peaks in known RAR$\alpha$ transcriptional targets *Rarb* (A) and *Abtb2* (B). (C-E) Representative RNAscope images (C) and CTCF quantification of *Sox2* and *Sox2ot* expression in LGE VZ (D), and their inverse correlation (E). (F-J) Representative images (F, G) and quantification of expression of Dll1 (H), Notch1 (I), NICD (J) in E14 control and *Foxc1* mutant LGE VZ from mice exposed to control or RA enriched diet. Means for each biological replicate (n = 3 embryos for each genotype-treatment) represented by colored shapes and corresponding technical replicates (n = 3 images per embryo) in grey circles, line and error bars shows mean and SD. *, P < 0.05, **, P < 0.01, ***, P < 0.001.
